# Dose-response relationship for the resistance of human insulin to degradation by insulin-degrading enzyme

**DOI:** 10.1101/2023.04.08.536135

**Authors:** Masaki Okumura, Tsubura Kuramochi, Yuxi Lin, Ran Furukawa, Kenji Mizutani, Takeshi Yokoyama, Mingeun Kim, Mi Hee Lim, Hyon-Seung Yi, Kenta Arai, Hiroshi Yamaguchi, Hironobu Hojo, Michio Iwaoka, Yoshikazu Tanaka, Sam-Yong Park, Kenji Inaba, Shingo Kanemura, Young-Ho Lee

**Author notes:** Corresponding authors: Masaki Okumura Young-Ho Lee.

## Abstract

Deeper understanding of the mechanism of the action of insulin and insulin-degrading enzyme (IDE) is a central theme in research into physiology and the pathophysiology of type 2 diabetes mellitus. Despite significant progress regarding the substrate recruitment, unfolding, digestion, and release by IDE, the structure and function of the insulin hexamer during the degradation cycle of IDE remain to be fully characterized. In the present study, we have characterized the behavior of human insulin hexamer in the absence of zinc. Using cryo-electron microscopy, we also observed that these hexamers represented a structure similar to that of T_6_ insulin. More interestingly, we also observed complexes in which some of their monomeric insulin components are partially distorted at their hexametric symmetry. This ensures that insulin determines the kinetics of its degradation by IDE without the requirement for zinc. These findings provide new information regarding the molecular events in insulin assembly and disassembly that permit its selective digestion by IDE.

## Introduction

Human insulin is a peptide hormone consisting of 51 amino acids that is produced by the β-cells of the pancreatic islets. The effects of insulin require the activation of insulin signaling following the binding of ligand to the insulin receptor, which can be characterized by a dose-response curve for the effect of insulin on circulating glucose concentration^1^. Insulin self-assembles into dimers, three of which combine with two zinc ions to form a 3-fold symmetrical hexamer^2, 3^. X-ray crystallographic and spectroscopic studies have demonstrated that the allosteric binding of hexameric insulin can be observed and the hexamer can be present in states that differ according to whether the residual numbers 1 to 6 of the B-chains of monomers exist in a tense (T) or relaxed (R) conformation. A conformational change is induced by the binding of phenol and chloride ligands to switch between R_6_ (a closed state)^4^, T_3_R_3_ (a semi-open state)^5^, and T_6_ (an open state) (Fig. 1a)^3^. Under physiological conditions, insulin is stored as zinc-induced hexamers, which form microcrystals within secretory granules, and ultimately zinc-free monomers are released from β-cell into the blood^6^. In the absence of zinc, at 0.2, 1, and 1.5 mM, the self-association statuses of human insulin are 50%, 75%, and 100% hexamer, respectively^7, 8^. A previous study also showed that the proportion of the insulin molecules that exist in the dimer form increases from 18% to 32% as the insulin concentration is increased from 0.05 to 1.0 mg/ml at a neutral pH and in the absence of zinc^9^. Thus, knowledge of the structure of insulin hexamers is limited to that obtained from crystallographic analyses of zinc-bound insulin, and it remains a great challenge to obtain structural information regarding the complexes formed between insulin molecules in the absence of zinc because of the dynamic and/or transient nature of their interactions.

**Figure 1.**
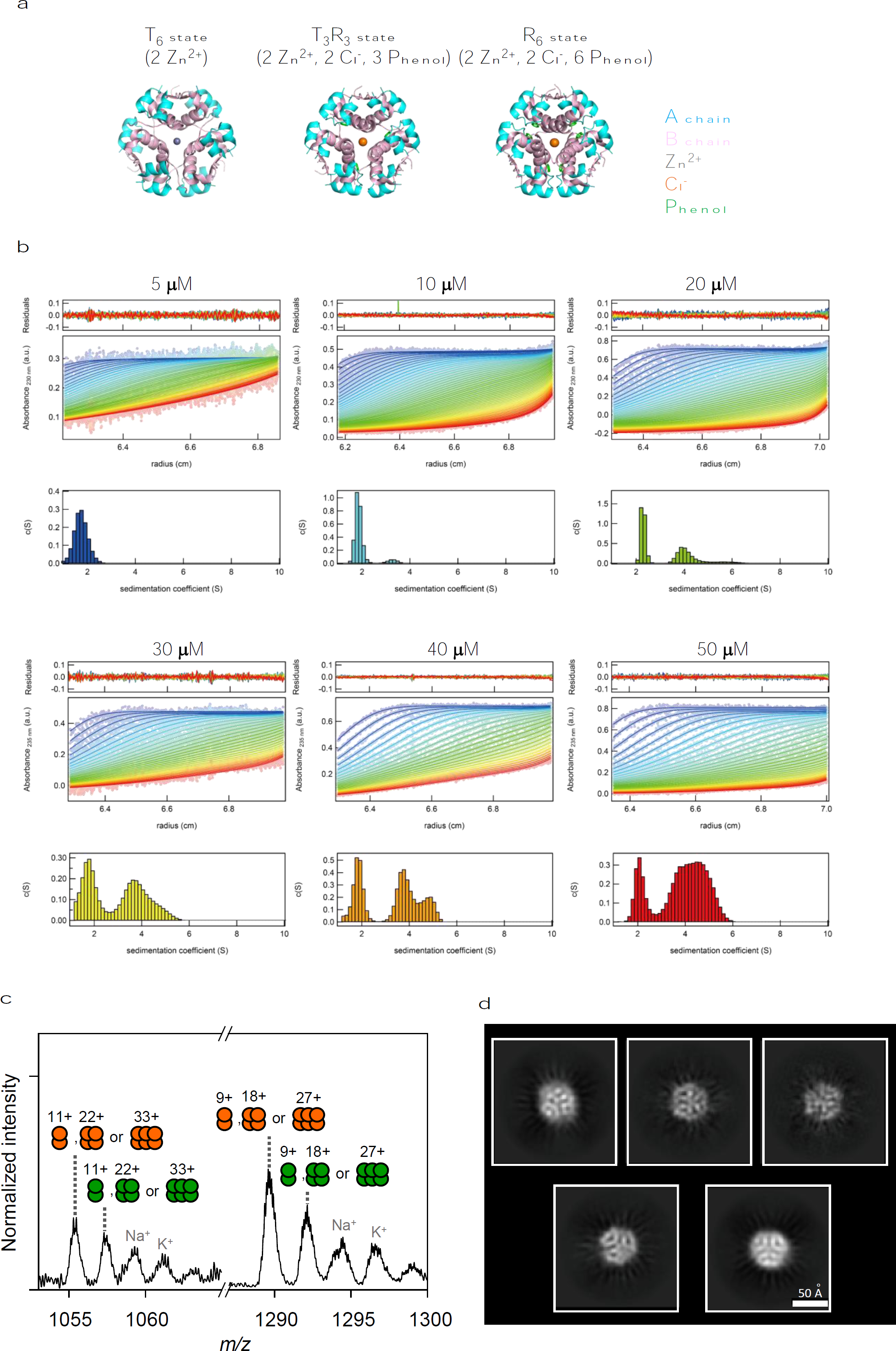
Dose-dependent hexamerization of human insulin in the absence of zinc. (a) Crystal structures of insulin hexamer T_6_ (PDB: 1MSO), T_3_R_3_ (1MPJ), and R_6_ (1ZNJ). (b) Apparent sedimentation coefficient distribution of human insulin at each protein concentration (5-50 μM). The data were obtained at 220 nm. (c) ESI-MS spectrum for human insulin at 50 μM. (d) Representative 2D class averages of single particle cryo-EM analysis.

Insulin-degrading enzyme (IDE) is ubiquitously expressed in mammalian tissues, including in the blood, skeletal muscle, liver, and brain^10, 11^, and is a zinc metalloprotease that hydrolyzes a broad range of bioactive peptides, such as insulin and amyloid β, which have diverse sequences and structures, and thereby regulate the amounts of these peptides present *in vivo*^12, 13^. Therefore, IDE-targeting therapeutics, such as inhibitors, are potential means of improving the regulation of blood glucose and treating type 2 diabetes mellitus (T2DM)^14, 15^. Extensive studies performed by Tang’s group have provided a molecular basis for the substrate recognition, unfolding, and degradation by IDE^16, 17^. The first published crystal structure of IDE revealed its planar structure, and its overall structure can be divided into two components, N- and C-clusters, which are connected by a short linker. IDE-N and IDE-C join together to form an encapsulated catalytic chamber to bind substrate^16–22^. The capture of substrate molecules involves at least two conformations during the catalytic cycle of IDE: it exists in an open conformation in the absence of substrate, and in a closed conformation when capturing substrate. Within the catalytic chamber, the interior surface of IDE is composed of specific substrate recognition sites: a catalytic cleft that coordinates a zinc ion, and an exosite that anchors the N-terminus of the substrate. The size of the catalytic chamber (∼13,000 A^3^) prevents IDE from encapsulating peptides of more than ∼80 amino acids in size. In addition to its proteolytic function, IDE suppresses the expression level of 140-residue long α-synuclein in β-cells in a nonproteolytic manner^23^, resulting in functioning as a chaperone. Despite significant progress in knowledge regarding substrate recruitment, unfolding, digestion, and release by IDE, the structure and function of the insulin hexamer during the catalytic cycle of IDE remain to be fully characterized. In the study presented here, we have demonstrated the characteristic behavior of the human insulin hexamer in the absence of zinc. Importantly, this is associated with resistance against degradation by IDE. Thus, the present study provides insight into the mechanism whereby the oligomerization of human insulin protects against its degradation by IDE in the absence of zinc.

## Results and Discussion

To characterize the oligomeric states of human insulin in the absence of zinc, the intrinsic viscosities and hydrodynamic radii were estimated using a Viscosizer TD (Malvern Panalytical). The *R*_h_, hydrodynamic radius, values for human insulin were increased dose-dependently between 5 μM and 80 μM (data not shown). When the crystal structures of human insulin were taken into the consideration, we found the *R*_h_ to be 1.50 nm as a monomer, 1.93 nm as a dimer, and 2.56 nm as a hexamer. To separate the oligomeric states, we next measured the sedimentation coefficient using analytical ultracentrifugation (AUC) at various concentrations. At concentrations <10 μM, almost all the single histograms had sedimentation coefficients of ≤1 S (Fig. 1b), which correspond to the monomer, based on the findings of the hydrodynamic radius analysis. Of interest, a population at a sedimentation coefficient of ∼4 S was also identified with the 50 μM concentration of insulin. When the distributions of the oligomeric states were plotted for each concentration between 5 μM and 50 μM, we found that the higher oligomeric state was more abundant because of a decrease in the number of monomers, despite the absence of zinc (Fig. 1b).

Subsequently, we investigated the oligomeric properties of human insulin at 50 μM using ESI-MS, and found that there were several oligomeric components, including hexameric insulin (Fig. 1c). Furthermore, we examined single particle cryo-electron microscopy of oligomeric insulin complexes. Image processing allowed us to visualize converged 2D class average images, showing stable structure with a diameter of about 50 Å, which corresponds to a hexametric complex similar to T_6_ oligomer (Fig. 1d). In addition, through visual inspection of cryo-EM images, we observed hexametric complex which is partially deformed. This suggests that interactions within hexamers in the absence of zinc are weaker than in the presence of zinc, rendering the structures more dynamic.

With respect to the regulation of circulating insulin concentration, IDE has been reported to bind insulin with high affinity (10 nM) and to rapidly cleave it (K_cat_ = 0.5–2 s^−1^) into peptide fragments^18, 24^, leading to its inactivation (Fig. 2a). Therefore, deeper understanding of the mechanisms of the effects of both insulin and IDE are central themes in research in physiology and into the pathophysiology of T2DM. Although the rate of bradykinin degradation by IDE is at least 20-fold higher than that of insulin, the mechanism involved remains largely enigmatic^25, 26^. To investigate the mechanism whereby insulin is degraded by IDE, we assessed the degradation of human insulin using purified recombinant human IDE. We incubated various concentrations of human insulin with 150 nM IDE and identified the peptide fragments generated using reverse-phase HPLC. At a concentration of 5 μM, insulin was completely digested by IDE within 20 min, but intact insulin remained when concentrations of 10 to 50 μM were used (Fig. 2b). Furthermore, the degradation rate constants decreased as the concentrations of human insulin were increased (Fig. 2c). Thus, whereas bradykinin degradation can be explained using the Michaelis-Menten equation^19^, it seems that insulin degradation by IDE is inhibited in a dose-dependent manner. The substrate size and degradative activity of IDE may be accounted for by the swinging door motion between IDE-N and IDE-C^19^, as follows. Briefly, when the door-swinging motion creates an 11 to 18 Å opening (Fig. 2a), this is sufficient for small peptides, such as bradykinin, to enter the catalytic chamber, while those of larger sizes, such as insulin monomer, are excluded. Besides providing information regarding the binding of monomeric insulin, we have also demonstrated that the equilibrium between the monomeric and hexameric forms of human insulin that lacks zinc determines the degradative activity of IDE, presumably because the insulin hexamer cannot access the active site of the enzyme.

**Figure 2.**
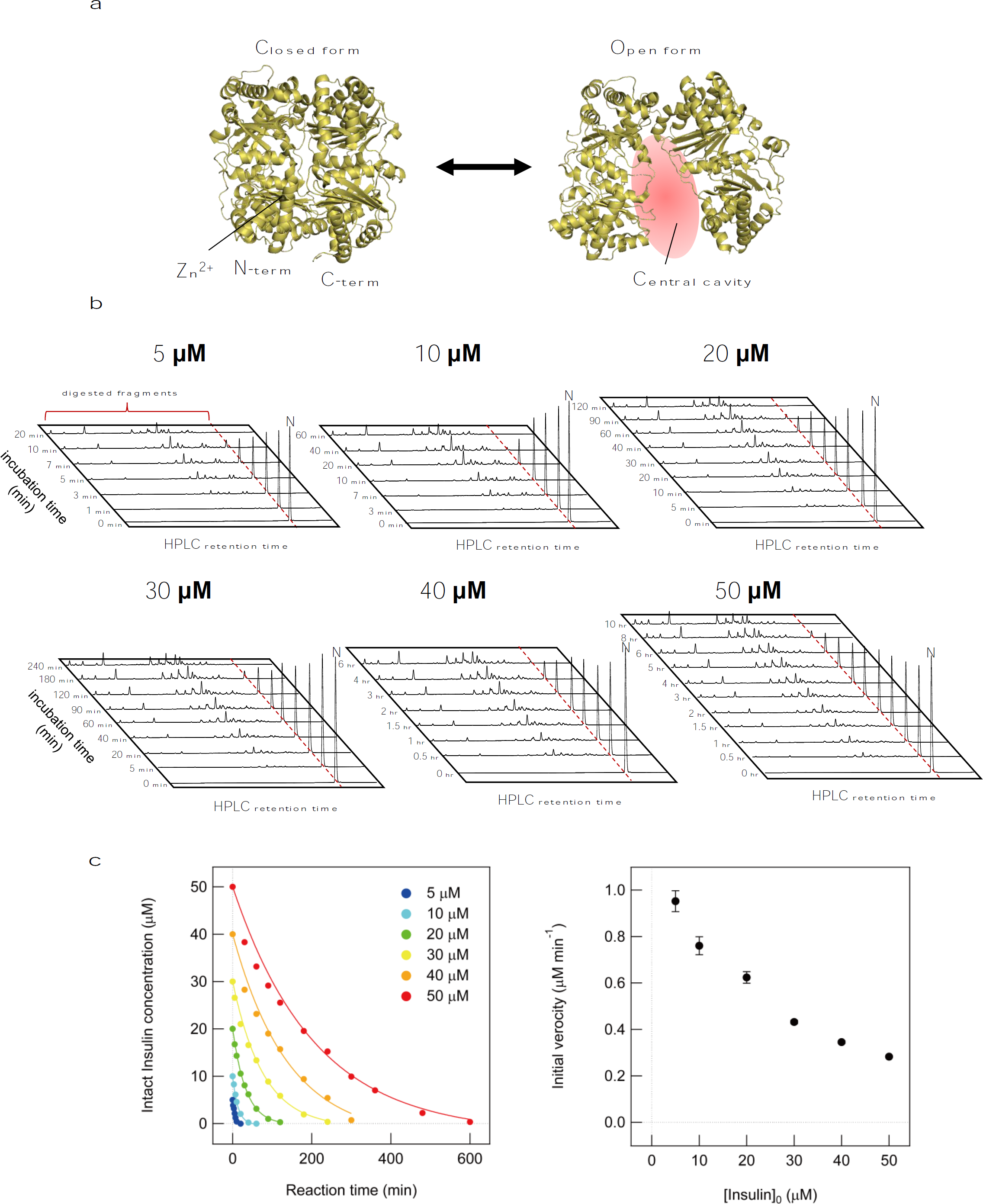
Dose-response relationship for the degradation of human insulin by IDE. (a) Crystal structures of closed (2JG4) and open (6B7Y) IDEs. (b) RP-HPLC profiles obtained during the digestion of each concentration of human insulin by IDE. (c) Kinetic profile of the enzymatic digestion of human insulin by IDE. Bars show mean ± SEM (n=3). (d) The representative 2D class averages of insulin hexametric complexes in the condition without zinc.

We next evaluated the effects of insulin degradation by IDE on insulin signaling in cultured cells (Fig. 3). Upon the stimulation of AML12 cells by human insulin, the phosphorylation of AKT was significantly increased. However, IDE inhibited this phosphorylation, presumably because of the degradation of human insulin by IDE. These data indicate that IDE likely regulates the physiological activity of human insulin through the binding of the monomer to the insulin receptor.

**Figure 3.**
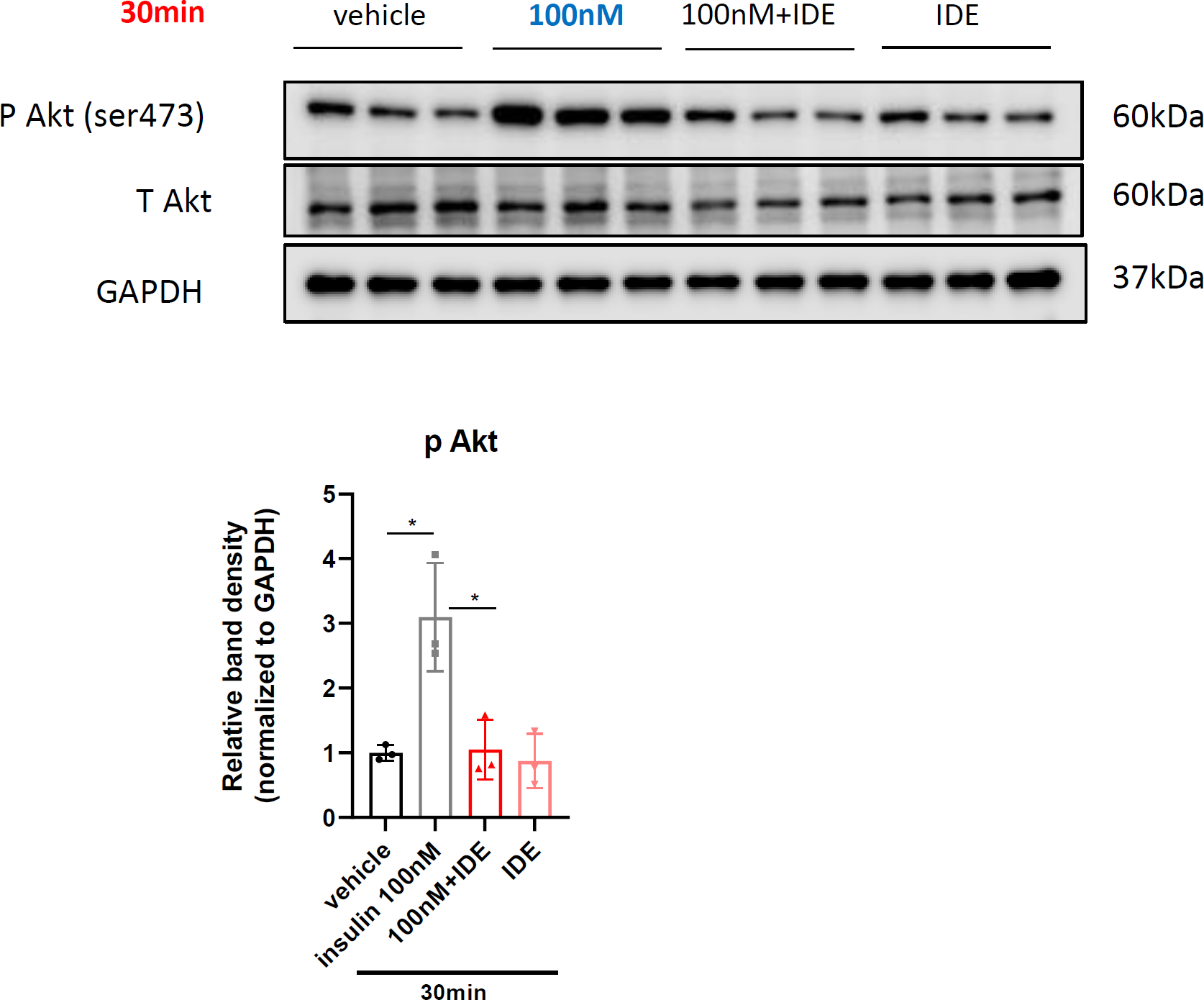
IDE reduces insulin signaling. AML12 cells were stimulated with 100 nM human insulin, IDE, or a mixture of these. Upper panel: results of the immunoblotting of whole-cell lysates. Lower panel: semi-quantitative results of immunoblotting. Bars show mean ± SEM (n=3). The signal intensities of phosphorylated AKT (Ser473) bands were normalized to those of AKT.

In summary, we have described the similarity in the structure of the insulin hexamer to T_6_ and the existence of hexamers in which some dimer units are partially dissociated, despite the absence of zinc. These findings highlight the resistance of the insulin hexamer to degradation by IDE. Our model of the method of insulin degradation by IDE provides a framework for the design of an insulin hexamer that resists degradation by IDE, which should accelerate the development of new treatment modalities for T2DM.

## Acknowledgments

This research was funded by a JSPS KAKENHI grant (grant numbers JP19K16092 (to SK) and 22H02205 (to MO)), a JSPS Grant-in-Aid for Transformative Research Areas (B) (grant number JP21H05095 (to MO)), the Japan Science and Technology Agency FOREST Program (grant number JPMJFR201F (to MO)), the Takeda Science Foundation (to KI and MO), the Mochida Memorial Foundation for Medical and Pharmaceutical Research (to MO), the Naito Foundation (to MO), the Uehara Memorial Foundation (to MO), the Terumo Life Science Foundation (to MO), Nagase Science Technology Foundation (to KI), the National Research Foundation of Korea (NRF) grant funded by the Korean government (grant number NRF-2022R1A2C1011793 (to Y-HL) and grant number NRF-2022R1A3B1077319 (to MHL)), the National Research Council of Science & Technology (NST) grant funded by the Korean government (grant number CCL22061-100 (to Y-HL)), and the KBSI fund (grant numbers C320000, C330130, and C390000 (to Y-HL)).

## Author Contributions

MO, TK, RF, and KS performed experiments, including the degradation by IDE. TK and RF prepared the purified IDE under the supervision of MO, HY, and SK. MO and MK performed the AUC experiments under the supervision of S-YP, and Y-HL. The cryo-EM study was conducted by TK and TY under the supervision of MO and YT. LY, MK and MHL conducted ESI-MS measurements and data analyses. H-SY carried out cell experiments and analyzed results. KA, HY, HH, MI, YT, S-YP, KI, MO, and Y-HL contributed to the analysis and interpretation of all results. MO, TK, RF, KA, HH, MI, KI, SK, and Y-HL discussed the insulin degradation mechanism by IDE. All the authors discussed the results, critically read the manuscript, and approved its submission. MO and Y-HL and designed and supervised the current work.

## Materials and Methods

### Materials

Human native insulin expressed in yeast (#12878) and bradykinin peptide (#3004/5) were purchased from Nacarai Tesque Inc. (Japan) and Tocris Bioscience (UK), respectively.

### Expression and purification of the recombinant protein

A pProEx-IDE-wt (#99014) plasmid containing cloned human IDE (Met42–Leu1019) was purchased from Addgene (UK). The plasmid encoded a 6-histidine tag at the N-terminus of the protein. A plasmid containing an IDE E111Q mutant was constructed using a QuikChange site-directed mutagenesis kit (Agilent Technologies, USA) and the pProEx-IDE-wt plasmid. The wild-type and E111Q forms of IDE were then overexpressed in the *Escherichia coli* strain BL21(DE3) by the addition of 0.5 mM isopropyl-β-D-thiogalactopyranoside and overnight culture at 20°C^16^. The IDE-overexpressing cells were disrupted in buffer A (50 mM Tris-HCl pH 8.1 and 300 mM NaCl) containing 1 mM phenylmethylsulfonyl fluoride using a homogenizer (Sonics and Materials, USA). The homogenates were centrifuged, and the supernatants obtained were loaded onto Ni-NTA Sepharose columns (Qiagen, Hilden, Germany), which were then washed with buffer A containing 20 mM imidazole, and the IDE was eluted with buffer A containing 200 mM imidazole. The eluted samples were concentrated to 500 μL volumes using Amicon Ultra filter units (MWCO, 10,000; Millipore, USA) on Mono-Q 5/50 GL anion exchange columns (Cytiva, USA) pre-equilibrated with 50 mM Tris-HCl (pH 8.1), and eluted using a linear gradient of 0–500 mM NaCl. The samples were then purified by size-exclusion chromatography using a Superdex 200 Increase column (Cytiva) pre-equilibrated with buffer A. The purified samples were replaced with 20 mM sodium phosphate buffer (pH 7.5) containing 100 mM NaCl and 10% glycerol, and stored at −80°C.

### Degradation assay

Insulin at various concentrations (5, 10, 20, 30, 40, and 50 μM) was mixed with 150 nM IDE and incubated at 30°C in 20 mM sodium phosphate buffer (pH 7.5) containing 100 mM NaCl. The reaction was then quenched using 0.5 M HCl and the mixtures were analyzed using RP-HPLC (GL Science, Japan) and a COSMOSIL 5C18-AR-II column (Nacalai Tesque), and eluted using a linear gradient of CH_3_CN in 0.05% TFA (5% to 85%) at a rate of 1% min^−1^, with detection using an absorbance of 220 nm. The HPLC peak area for the native substrate was plotted against the degradation time, and the degradation rate constant for IDE was determined by curve-fitting using the following exponential equation:

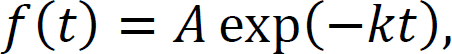

where A and k are the amplitude and degradation rate constant, respectively. All the calculations were performed in Igor Pro (WaveMetrics, USA).

### Assessment of insulin signaling in AML12 cells

AML12 cells were propagated in T-75 flasks and experiments were carried out in 6-well plates. AML 12 cells were seeded at a density of 3×10^5^ cells in each well and cultured in DMEM F-12 medium (Welgene, South Korea) containing 10% fetal bovine serum (Hyclone; Thermo Fisher Scientific, USA) and 1% penicillin and streptomycin (Welgene, South Korea). After overnight culture, the cells were treated with 10 nM or 100 nM human insulin in 20 mM sodium phosphate and 100 mM NaCl buffer ± 0.5 μM IDE at 30°C for 15 or 30 min.

### Western blot analysis

Primary hepatocytes and AML12 cells were washed with cold PBS and lysed in a TissueLyser II in a buffer (50 mM Tris-HCl, pH 7.4; 150 mM NaCl; 1 mM EDTA, pH 8.0; 0.1% Triton X-100) containing a protease inhibitor cocktail (#11836145001, Roche, Switzerland) and phosphatase inhibitors (#04906837001, Roche) on ice for 30 min. The lysates were then centrifuged for 15 min at 16,000 × *g* at 4°C and their protein concentrations were determined using a BCA Protein Assay Kit (#23227, Thermo Fisher Scientific, USA), according to the manufacturer’s instructions. Lysates containing 50 µg protein were separated by electrophoresis on 8–12% polyacrylamide gels and the separated proteins were then electrotransferred to 0.45 μm PVDF membranes (#IPVH00010, Millipore, USA) at 200 mA for 2 h. The membranes were blocked with 5% skimmed milk (#T145.2, Carl Roth, Germany) in TBS/T buffer (20 mM Tris-HCl, 150 mM NaCl, 0.1% Tween 20, pH 7.6) for 1 h and then incubated with primary antibodies (anti-phospho-AKT-Ser473 (#4060, Cell Signaling Technology, USA), anti-AKT (#9272, Cell Signaling Technology), or anti-GAPDH (#2118, Cell Signaling Technology)) overnight at 4°C. The membranes were then washed three times with TBS/T and incubated with secondary antibody (Goat Anti-Rabbit IgG (H+L)-HRP Conjugate, #1706515, Bio-rad, USA) for 1 h at room temperature. They were then scanned with an Odyssey Western Blot Scanner (LI-COR) and analyzed using Image Studio Lite (LI-COR).

### Analytical ultracentrifugation experiments

Insulin at various concentrations in 20 mM sodium phosphate buffer (pH 7.5) containing 100 mM NaCl was loaded into the measurement cell in 400 μL volumes, and 420 μL volume of 20 mM sodium phosphate buffer (pH 7.5) containing 100 mM NaCl was loaded into the reference cell. Each insulin sample solution was first centrifuged at 645×g, and sedimentation velocity measurements were made from 261,668×g and at 37°C. The obtained absorbance data were analyzed with sedfit software using solvent parameters (partial specific volume, density, and viscosity) calculated using sednterp.

### Electrospray ionization–mass spectrometry

A solution of human insulin (50 μM) in 20 mM ammonium acetate (pH 7.5) was prepared. The capillary and nozzle voltages were set to 5.8 and 2 kV, respectively; and the gas temperature, drying gas flow rate, and nebulizer pressure were 300°C, 12 L/min, and 60 psig. More than 500 spectra per sample were obtained and mean values were calculated. The capillary voltage was set to 5.8 kV and the nozzle voltage was 2 kV. The gas temperature, drying gas flow rate, and nebulizer pressure were 300°C, 12 L/min, and 60 psig, respectively.

### Cryo-EM grid preparation, data collection, and image processing

3 μl of 50 μM purified insulin in 20 mM sodium phosphate buffer containing (pH 7.5) containing 100 mM NaCl was applied onto a grow discharged Quantifoil R1.2/1.3 Cu 200 grid (Quantifoil Micro Tool GmbH) and subsequently blotted by filter papers and plunged into liquid ethane for the vitrification using a Vitrobot Mark IV (Thermo Fisher Scientific). The vitrified specimen was observed with a CRYO ARM 300 II transmission electron microscope (JEOL) equipped with a cold field emission gun operated at 300 kV accelerating voltage. Insulin complex images were recorded with a K3 camera (Gatan) as movie micrographs at the total electron exposure on the specimen was around 80e-Å^−2^ and the nominal magnification at 60,000. The automated data collection was performed with the SerialEM program^27^. The image processing was performed with the CryoSPARC program^28^. Briefly, 6,699 movie micrographs were motion-corrected. Insulin complex images were picked with the blob picker implemented in the CryoSPARC. The total 6,170,486 particles were sorted into homogeneous subgroups by 2D classification. The representative 2D class averages are shown in Figure 1d.

